# Skipping and sliding to optimize target search on protein-bound DNA and RNA

**DOI:** 10.1101/2020.06.04.133629

**Authors:** Misha Klein, Tao Ju Cui, Ian MacRae, Chirlmin Joo, Martin Depken

**Affiliations:** Kavli Institute of Nanoscience and Department of Bionanoscience, Delft University of Technology, Delft, The Netherlands; Department of Integrative Structural and Computational Biology, The Scripps Research Institute, La Jolla, United States

## Abstract

Rapidly finding a specific nucleic-acid sequences in a large pool of competing off-targets is a fundamental challenge overcome by all living systems. To optimize the search and beat the diffusion limit, it is known that searchers should spend time sliding along the nucleic-acid substrate. Still, such sliding generally has to contend with high levels of molecular crowding on the substrate, and it remains unclear what effect this has on optimal search strategies. Using mechanistic modelling informed by single-molecule data, we show how sliding combined with correlated short-ranged skips allow searchers to maintain search speed on densely crowded substrates. We determine the conditions of optimal search, which show that an optimized searchers always spend more than half its time skipping and sliding along the substrate. Applying our theory to single-molecule data, we determine that both human and bacterial Argonaute proteins alternate between sliding 10 nt and skipping 30 nt along the substrate. We show that this combination of skipping and sliding lengths allows the searcher to maintain search speeds largely unaffected by molecular roadblocks covering up to 70% of the substrate. Our novel combination of experimental and theoretical approach could also help elucidate how other systems ensure rapid search in crowded environments.

---

The flows of information in the central dogma of molecular biology rely on proteins efficiently finding specific DNA and RNA sequences. The sheer size of intracellular and nuclear volumes restrict the rate of finding a target through diffusive collisions alone^1–4^. Still, measured rates exceed this theoretical upper bound by up to two orders of magnitude^5^. A substantial body of both theoretical^1,6–16^ and experimental^17–31^ research has established that proteins can utilize a combination of 3D diffusion through solution and 1D lateral diffusion along the substrates to facilitate the search process and reach the measured search speeds.

Most theoretical and *in vitro* biophysical studies have focused on search along bare and double-stranded DNA, which is simple in structure and presents almost no physical obstructions to sliding. However, *in vivo* DNA and RNA are highly decorated with DNA and RNA binding proteins and secondary structures^11,15,32–35^. While searching along the substrate, the protein stays in close proximity at all times, and benefits of 1D motion will be limited without a specific mechanism for bypassing roadblocks. The motion is typically split up into 1-nucleotide (nt) sliding steps, where each base is interrogated by the searcher, and various forms of base skipping, where bases are not interrogated by the searcher. The skipping behavior falls into three broad classes: *hops/micro-dissociations*^1^ (**Supplementary Figure 1a**), where bases are skipped due to either partial dissociation, where the searcher remains within the electrostatic-interaction range but is disengaged from the hydrogen bonds furnishing the sequence dependence (**Supplementary Figure 1a** i), or the protein has two binding pockets, allowing it to perform the walk shown in **Supplementary Figure 1a ii**; *intersegmental transfers*^6,8–10,36^ (**Supplementary Figure 1c**), where the substrate flexibility enables two distal sites on the substrate to come into close proximity in space, allowing a searcher to directly transfer from one substrate segment to another; *distinct protein conformations* (**Supplementary Figure 1d**), where the protein is be able to adopt one conformation allowing for stronger sequence recognition, and another for faster 1D diffusion^7^.

Independent of the mechanism through which base skipping occurs, their frequency and length will influence the average time needed for the protein to locate the target sequence. To construct a general experimental assay for characterizing the distribution of skipping distances, we here introduce a stochastic skip-n-slide (sNs) model that accounts for sliding steps, skips of any form, and 3D diffusion. We provide the general conditions for optimal search, and show that there are two locally optimal search strategies: one utilizing sliding but not skipping, and one with full sNs-search. A sNs-search strategy allows the searcher to mitigate the effects of roadblocks, and we establish analytical conditions for when search speeds remain high in the face of molecular crowding along the DNA.

In a recent experimental study^32^, we showed that Argonaute (Ago)—a searcher along single-stranded (ss) nucleic-acid substrates in eukaryotic post-transcriptional regulation^37–39^ and prokaryotic host defense^40,41^ — can bypass protein and secondary-structures without dissociating. These findings hint at how Argonaute can perform efficient facilitated diffusion on molecularly/structurally crowded substrates. By applying our theory to single-molecule Förster resonance energy transfer (smFRET) experiments, we show that both human Argonaute 2 (hAgo2) and *Clostridium Butyricum* Ago (CbAgo) use sNs-search. On an unstructured substrate, the searchers slides a distance of around 10 nt before being interrupted by short-range intersegmental transfers where about 30 nt are skipped. We end by showing that Ago’s search is consistent with sNs optimal search, and can progress essentially unhindered by roadblocks for up to 70% coverage of the substrate.

## Results

### Detecting skip-N-slide target search

Previous experimental approaches to characterizing diffusive motion rely on tracking the molecule’s movement across hundreds of nucleotides in order to span the diffraction limit^17,18,21,22,25,27,28,30,31^. As lateral excursions might not reach such lengths^42^, we seek a method that is capable of detecting shorter excursions. Following earlier experimental work^19,29,32,43,44^ we consider experimental designs with a substrate containing two specific binding sites (traps), separated by non-specific sites. With the searcher trapped long enough to be detected, this design allows us to accurately determine the shuttling time between traps. We begin by showing that the shuttling time from one trap to the other depends on the number of non-specific sites in between them. We base our argument on the assumption that 1D diffusion can be seen as consisting of repeated sNs cycles. Each cycle is composed of approximately 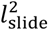 diffusive single-nucleotide steps, that typically scan a segment of length *l*_slide_, followed by a multi-nucleotide skip (**Fig. 1a**) of average distance *μ*_skip_ in either direction, spread over a distance *σ*_skip_ (**Fig. 1b**). After a hop/micro dissociation^1^ event for a searcher with only one binding pocket, the searcher is most likely to rebind where it unbound from, and such events are described by *σ*_skip_ > *μ*_skip_ = 0. Both hopping with multiple binding modes and intersegmental transfers^6,10^ are likely to rebind some appreciable distance away from previous binding site, and such events are described by *μ*_skip_ > *σ*_skip_ > 0. In **Figs. 1b** and **d** we illustrate how this last situation leads to a central gap in the distribution of nucleotides covered in one round of sliding and skipping.

**Fig. 1.**
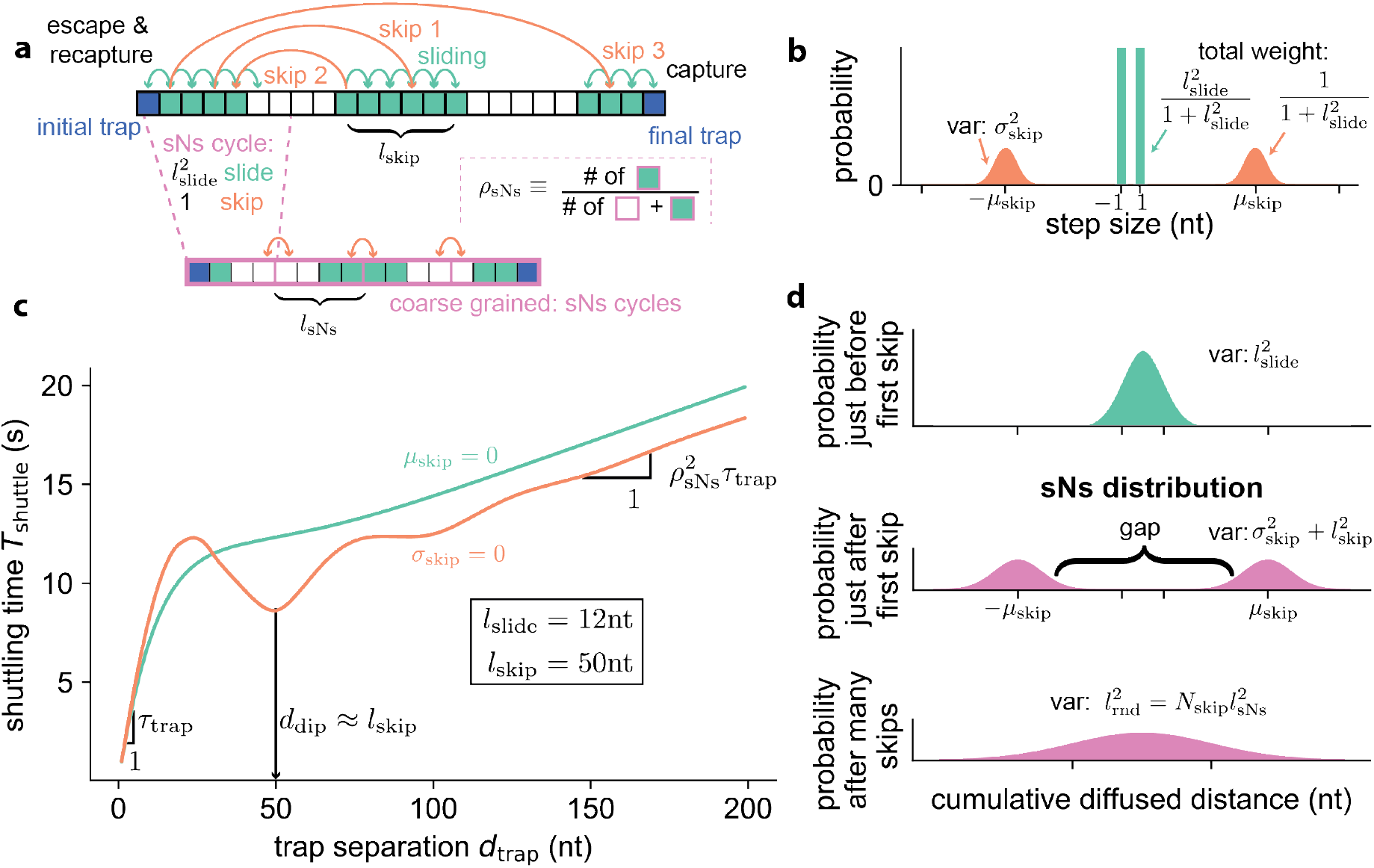
A tandem trap assay to characterize sNs-search. **a,** (top) Schematic of a shuttling event. Starting from the leftmost trap (traps indicated in blue), the protein uses a combination of single-nucleotide steps (sliding, turquoise) and larger steps (skipping, orange) to reach the opposite trap, after possibly getting recaptured at the initial trap several times. (bottom) After multiple skips, interspersed by 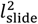 sliding steps, a coarse-grained description of the shuttling process is used to describe long distance shuttles. Within this coarse-grained description, the protein skips between neighboring sites that are all the size of a typical sNs cycle (*l*_sNs_) (pink boxes), and all have the same fixed scanning density (*ρ*_sNs_) (given by the fraction of turquoise to white in the pink boxes). **b,** Single-step distribution of the sNs model. The protein either slides to a neighboring site (turquoise bars) or skips to sites located a distance *μ*_skip_ ± *σ*_skip_ away (orange peaks). **c,** Representative numerical solutions (**Supplementary Information**) for shuttling time versus trapping distance. In both cases shown, the mean-square distances 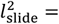 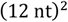 and 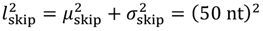 are the same. The composition of the skipping distance differs in the two cases though, with the orange curve corresponding to precise skips (*l*_skip_ = *μ*_skip_ and *σ*_skip_ = 0) and the turquoise curve corresponding to skips with maximal spread (*l*_skip_ = *σ*_skip_ and *μ*_skip_ = 0). Consequently, the orange and turquoise curves correspond to a gapped and non-gapped sNs distribution respectively (see middle panel of **d**). **d,** Distribution of searcher position conditioned on skips. (top panel) The protein covers a mean-square distance 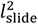 between skips. (middle panel) The first skip takes the protein *μ*_skip_ away in either direction, with a variance 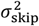 at either landing site (the sNs distribution). (bottom panel) *N*_skip_ repeated sNs cycles result in a distribution with variance 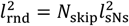 equal to the typical distance traversed before unbinding into solution.

### Linear increase in shuttling time at small trap separations

For rapid sliding without skipping, we recently showed^32^ that the total shuttling time between two traps is given by

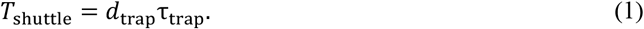

Here τ_trap_ is the average escape time from the trapping sequence, and *d*_trap_ is the distance between traps. Even when skipping is allowed, we can bring traps close enough that there is no time to skip before shuttling to the other trap (*d*_trap_ < *l*_slide_, see **Fig. 1c**). We can therefore determine τ_trap_ by fitting **Equation 1** to measured shuttling times at small trap separations.

### Linear increase in shuttling time at large trap separations

If traps are so far apart that there are many sNs cycles in a single shuttling event, then large-scale dynamics will resemble that of a simple random walk with a mean-squared step length (**Figs. 1a, d**)

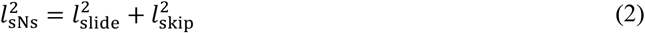

per sNs cycle. Here we have introduced the mean-square skipping distance 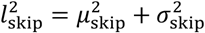. In the **Supplementary Information** we use a description conditioned on skipping events (**Fig. 1d**) to construct (and numerically validate, **Supplementary Figure 2**) a scaling argument that shows that we should expect the shuttle time to grow linearly (**Fig. 1c**) as

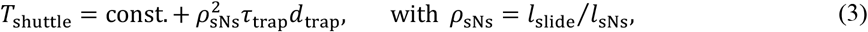

for large trap separations. Here we have introduced the scanning density *ρ*_sNs_ as the fraction of unique bases interrogated by the searcher over a single sNs cycle (dashed box in **Fig. 1a**). By fitting both the initial and the final slope of measured shuttling time vs. trap separation, we can now determine the scanning density of the searcher as well as the trapping time.

### Shuttling times can display a minimum at intermediate trap separations

In between the two aforementioned linear regimes, the shuttling time can vary non-monotonically and produce a local minimum. To see this, consider the distribution of distances traveled in a single sNs cycle (**Fig. 1d**, middle panel), which we shall refer to as the sNs distribution. There is a central gap in the sNs distribution when the average mean-squared skipping distance 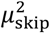 is substantially greater than the total mean-squared spread 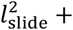 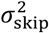 around the landing positions (this situation is illustrated in the middle panel of **Fig. 1d**). For trap separations that lie well within the gap, skipping is not beneficial, and the searcher must slide across. The local minimum in shuttling times appears when the second trap sits precisely one skipping distance away from the first trap (*d*_trap_ = *μ*_skip_), as this allows the searcher to skip straight into the second trap with little additional sliding needed. For there to be a substantial gap in the sNs distribution, we need 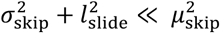, implying that *l*_sNs_ ≈ *μ*_skip_ ≈ *l*_skip_ (see **Equation 2** and accompanying text). That is, if a clear dip is observed in the shuttling time vs. trap separation, its position reports the average skipping distance (*d*_dip_ = *μ*_skip_, **Fig. 1c**). Only intersegmental transfers and a two-binding pocket mechanism allow for a gapped distribution, and a local minimum in shuttling time vs. trap separation curve. For systems with only one substrate binding site, the former mechanism is ruled out, and any observed local minimum will be evidence of intersegmental transfers.

### Lateral diffusion of Argonaute

To demonstrate the power of the above theoretical analysis, we focus our proof-of-principle on hAgo2 and CbAgo, which respectively use a 22-nt guiding RNA to bind ssRNA^37–39^ and a 23 nt guiding DNA to bind ssDNA^45^. Ago potentially displays several variants of skipping (**Supplementary Figure 1**), but which one is dominant is not known. Electrostatic interactions between guide/protein and target allow for micro-dissociations. The search along flexible single-stranded substrates heightens the possibility of short-ranged skips via intersegmental transfer. hAgo2 is also known to undergo a conformational change during target site recognition^46^, which possibly takes Ago between search and recognition conformations^47^.

To probe lateral diffusion on length scales as short as a few nucleotides, the relevant scale of local target search by (guided) target searchers^32,42,43^, we here used smFRET (**Fig. 2a**). To trap diffusive excursions for long enough to detect it (>100 ms), and have it complete before photobleaching (<700s), we designed single-stranded thymine (CbAgo^32^) and uracil (hAgo2, **Supplementary Figure 3**) repeats that contain two 3-nt traps and two 4-nt traps respectively (**Fig. 2a**). To observe protein binding, one trap is labeled with an acceptor fluorophore (Cy5), while the guide (irreversibly anchored to the Ago protein) is labeled with the donor fluorophore (Cy3). High FRET efficiency is thus observed when the protein binds to the labelled trap, whereas lower FRET efficiency is obtained when Ago binds to the unlabeled trap (**Fig. 2b**, **Supplementary Figure 3**). This construct allows us to observe both binding to the substrate, as well as which trap Ago is bound to. To reduce the background fluorescence, traces were recorded using total internal reflection (TIRF) microscopy. Using TIRF allows us to focus only on 1D diffusion during a single binding event, as freely diffusing proteins move too fast across the evanescent field to be detected.

**Fig 2.**
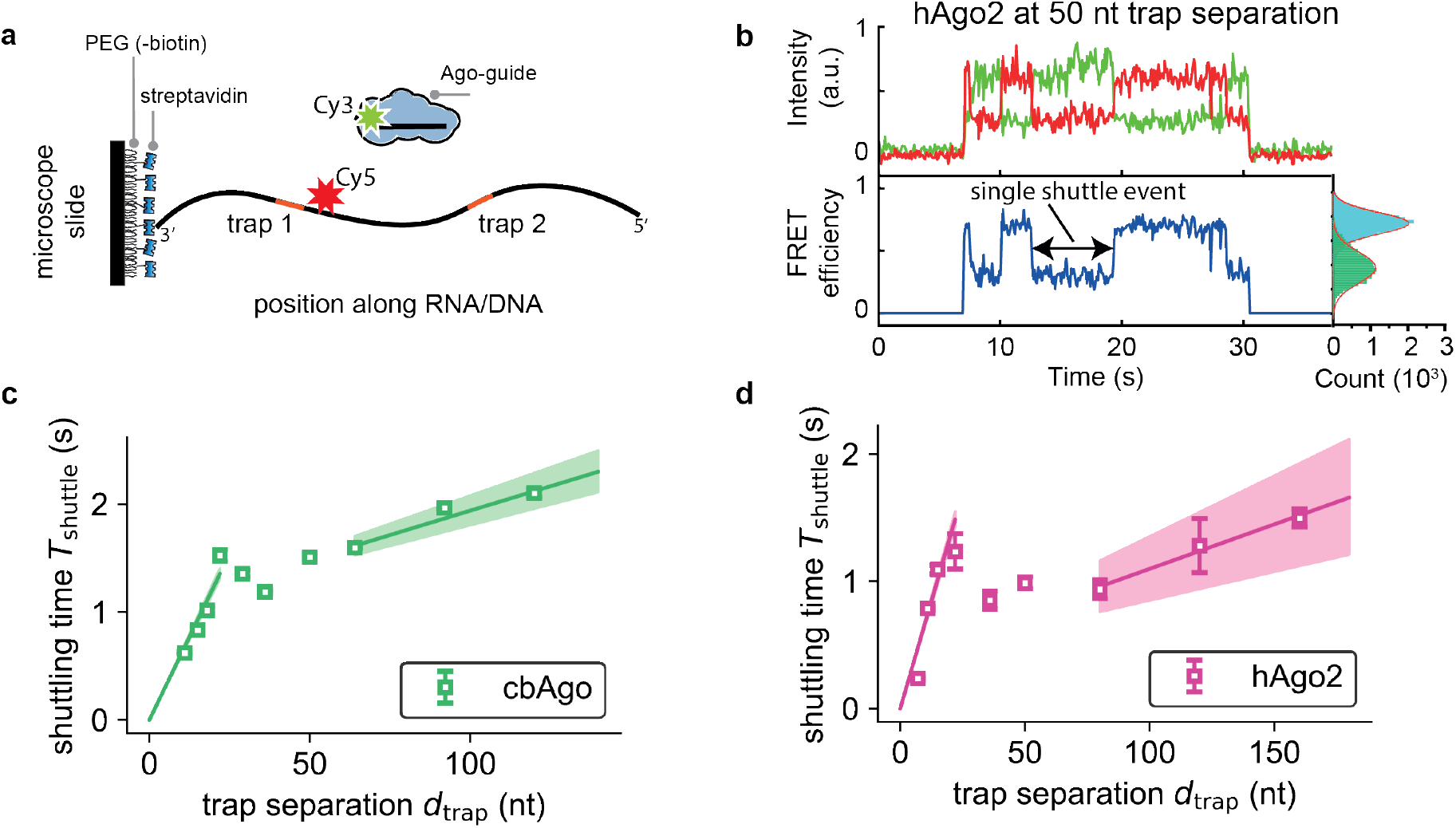
smFRET realization of tandem target assay using Argonaute. **a,** Schematic of our experimental assay. Single-stranded DNA/RNA constructs containing the two trapping sequences (shown in red) are labelled with the acceptor die and attached to the microscope slide via a 3’ biotin-streptavadin linker. The Ago-guide complex is labelled with the donor die. **b,** Representative trace for hAgo2 at a trap separation of 50nt. The top panel shows donor (green) and acceptor (red) signals, and the bottom panel shows the corresponding FRET efficiency, and the side panel shows histogram of all FRET efficiency values obtained for the population of molecules. **c,** Shuttling time vs. trap separation for CbAgo. Solid lines represent linear fits to data points (average ± sem) at 11 nt, 15 nt, 18 nt, 22 nt (initial slope) and 64 nt, 92 nt, 120 nt (final slope) **d,** Shuttling time vs. trap separation for hAgo2. Data points at 7 nt, 11nt, 15 nt (initial slope) and 80 nt, 120 nt, 160 nt (final slope) are used for linear fits. Shaded regions in **c** and **d** represent 95% confidence interval obtained using bootstrapping (**Supplementary Information**).

As shown in **Fig. 2b**, the FRET efficiency shifts almost instantaneously between the levels corresponding to the two trap locations. Though smFRET has a high spatial resolution, the total time spent diffusing between traps has fallen below our time resolution (30-100 ms). Still, we capture the time it takes to shuttle between traps at nucleotide resolution. This enables us to compare the experimental curves of **Figs. 2c** and **c** to the theoretical curves in **Fig. 1c.** As we have no evidence of Ago having the two substrate binding pockets, we take the appearance of a minimum in measured shuttling times vs. trap separation as evidence that Ago skips through intersegmental transfers.

### Argonaute skips three times as far as it slides

To estimate the trapping time *τ*_trap_ we fit **Equation 1** to the initial linear part in **Figs. 2c** and **d**, resulting in 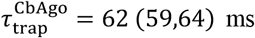 and 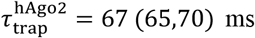 (numbers within brackets indicate 95% confidence interval, **Supplementary Information**). Next, we determine the scanning density of an sNs cycle by fitting **Equation 3** to the final linear part of **Figs. 2c** and **d**. The resulting scanning densities 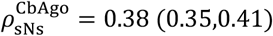 and 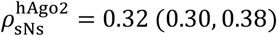 indicate that both Ago skip about three times the distance it slides. From our previous argument we know that the dip visible in the shuttling time in **Figs. 2c** and **d** essentially reports on the length of the skips, implying *l*_skip_ ≈ 30 nt. Using the measured scanning densities we also have *l*_slide_ ≈ 10 nt (**Equation 3**).

### Optimal sNs-search strategies

Skipping clearly helps the searcher to swiftly reach distant sequences, but could also lead to the target being missed if too few sliding steps are used in between skips. If too many sliding steps are used in between skips, the protein will repeatedly scan sequences, rendering the search inefficient. Here we establish the conditions under which sNs search reduces the time needed for protein to find a target within a genome or mRNA pool. To this end, we calculate the average time to find a target site through sNs search, interrupted by complete dissociation into solution and rebinding. We start by considering bare substrates, and later include the effects of roadblocks.

Our model (**Supplementary Information**) covers the scenario presented in **Fig. 3a**, where even though the target site is located between the binding and unbinding sites of a searcher during a 1D search round, it can still be skipped over and missed. We optimized the time needed to find a target amongst *L* sites,

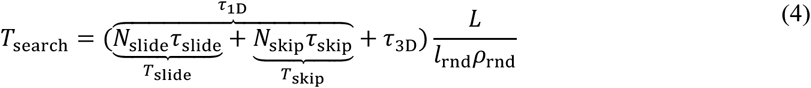

with respect to the typical number of skips (*N*_skip_) and sliding steps (*N*_slide_) performed in a single sNs cycle. In **Equation** 4, *T*_slide_ and *T*_skip_ denote the total time spent on sliding and skipping during every round of facilitated diffusion. These total times (*T*’s) are the product of the number of times the searcher slides or skips (*N*’s) and the time required to perform an individual step and interrogate the landing site (*τ*′*s*) (see **Supplementary Information** for details). To expand upon existing models^6,7,16^, we let the effective scanning density *ρ*_rnd_ denote the fraction of sites interrogated during one 1D lateral diffusion event (see dashed box in **Fig. 3a** and **Supplementary Figure 4**). The effective scanning density should be contrasted with the scanning density (*ρ*_sNs_), which gives the chance of an intervening base being checked in a single sNs cycle.

**Fig. 3.**
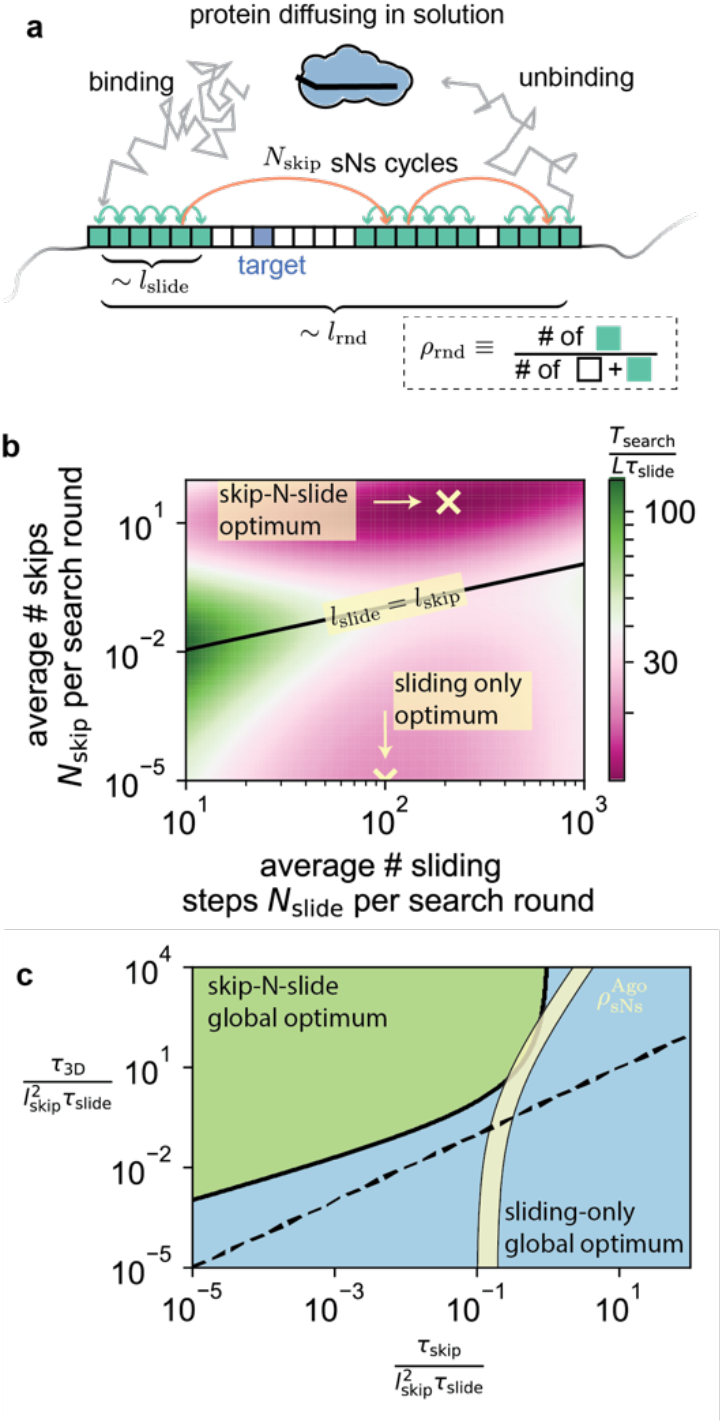
Optimal search times. **a,** Schematic single search round. In search of the target site, the protein binds, and cycles through an average of *N*_skip_ sNs cycles before it unbinds into solution. Only sites slid past (at least once) are interrogated (turquoise), resulting in a scanning density *ρ*_rnd_ between the binding and unbinding sites. **b,** Search time versus *N*_slide_ and *N*_skip_ for *l*_skip_ = 30 nt and *τ*_3D_ = 10*τ*_skip_ = 100*τ*_slide_. Region above the solid line represents sparse scanning (*ρ*_sNs_ < 0.5) while the region below the line represents dense scanning (*ρ*_sNs_ > 0.5). The sliding only optimum (on the boundary of zero skips) and the sNs optimum are indicated with crosses. **c,** Diagram showing when the sNs minimum is the global minimum 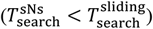 in green, as a function of the effective parameters that control the system (see **Supplementary Information** for details).

In the **Supplementary Information** we show that the search time has two local minima (**Fig. 3b**): one sliding-only minimum and one sNs minimum. We further show that in the high scanning-density region, where *l*_skip_ ≪ *l*_slide_, a minimum search time occurs on the boundary 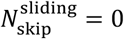. At this local minimum, the protein spends half its time diffusing through solution and the other half sliding (*τ*_1D_ = *τ*_3D_, see **Supplementary Information** for derivation), in agreement with the known minimal search time when *a priori* assuming that there are no skips^6,7^.

In the low-scanning density region, where *l*_skip_ ≫ *l*_slide_, the protein is at its fastest if it spends half its time skipping (*T*_skip_) and the other half on a combination of sliding (*T*_slide_) and diffusing through solution (*τ*_3D_) (see **Supplementary Information** for the general condition of optimality)

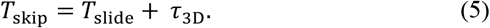

Consequentially, at optimum, the time spent diffusing laterally (*τ*_1D_ = *T*_skip_ + *T*_slide_) exceeds the time spent diffusing through solution (*τ*_1D_ > *τ*_3D_). This result is general, and remains true also outside the strictly low scanning-density regime (**Supplementary Information**).

As there are local minima in both the sparsely and densely scanned regions (**Fig. 3b**), the global optimal search strategy is defined by which of these two minima have the smallest search time. In **Fig. 3c** we identify the parameter values that make the sNs minimum the global minimum, and show the thin slice of (possibly only locally) optimal sNs systems compatible with the Ago in yellow (signifying scanning densities between 0.3 and 0.4). Next, we consider the implications of the three major types of skips shown in **Supplementary Figure 1** on our general model’s outcomes, and then turn to the effect of roadblocks.

### Skipping in the form of hopping/micro-dissociation

First, we consider skips resulting from hopping or micro-dissociations. The contribution of this search mode to the overall search time can be probed by altering the salt concentration^12,17,21,28,48^. At increasing salt concentrations, more counter ions need to be displaced when completing a skip, and more time is needed (**Supplementary Figures 1a,b**). It is therefore expected that the effective skipping length (*l*_skip_) increases with increasing salt concentration. Raising *l*_skip_ will lower the scanning density *ρ*_sNs_, which could be observed as an outward shift of the local minimum in the shuttling time vs. trap separation curve, accompanied by a decrease in its terminal slope (**Supplementary Figure 5**).

### Skipping in the form of intersegmental transfers

Next, If skips are the result of intersegmental transfers, their contribution can be probed by applying pulling forces to the substrate^36^. Tension will suppress large-scale looping back of the substrate (**Supplementary Figures 1a** and **c**), and distant sites will grow less likely to be close enough to allow a skip. Therefore we expect *l*_skip_ to decrease with tension, as would be evidenced by an increase in the terminal slope of the shuttling time vs. Trap curve (**Supplementary Figure S5**). Note that for single-binding pocket searchers, observing a minimum in the shuttling time vs. trap separation curve directly implies that intersegmental transfers are utilized (see above).

### Skipping to overcome the search-speed/stability paradox

Lastly, it was previously pointed out that strong sequence recognition suppresses lateral diffusion, leading to an apparent search-speed/stability paradox at physiological conditions^7^. It was also suggested that this paradox could be circumvented if the protein is able to switch between two conformations: one with strong sequence specificity/recognition and low diffusion constant *D*_rec_, and one with little sequence specificity and high diffusion constant *D*_srch_ (**Supplementary Figure 1d**). In the context of sNs-search, the non-specific fast mode can be seen as a diffusive skip 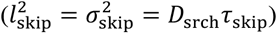, while the specific slow mode corresponds to sliding. Interestingly, in the low scanning density regime, the optimal search speed is dictated by the (large) diffusion constant of the search mode, and not the (small) diffusion constant in the recognition mode (**Supplementary Information**).

How is recognition still ensured when diffusion rates are set by the sequence insensitive search mode? In **Supplementary Figure 4** we show the effective scanning density *ρ*_rnd_ as dependent on the scanning density *ρ*_sNs_. From this figure it is evident that the effective scanning density can be close to one, even when the scanning density in a single sNs cycle is small. Therefore, the frequent use of skips offers a direct solution to the search-stability paradox introduced by Slutsky and Mirny^7^, as our calculation identifies the low scanning-density regime as a regime where the diffusive speed is set by the fast mode, while essentially all bases traversed during a 1D excursion are still checked.

### Search on molecularly crowded substrates

We close by asking to what extent search times change when the DNA and RNA is covered with roadblocks, such as other proteins or secondary structures^11,14,15,33–35^. If the target searcher is unable to displace roadblocks, the obstacles can be surpassed only through unbinding followed by rebinding (case I in **Fig. 4**), or through a skip (case II in **Fig. 4**). In case I, where unbinding is used to bypass roadblocks, the only requirement for efficient search is that the distance between roadblocks (*l*_bare_) should exceed the typical distance covered during 1D lateral diffusion (*l*_rnd_ < *l*_bare_). In case II, where skipping is used to bypass roadblocks, the same argument says that efficient search is possible only when sliding is not hindered by roadblocks (*l*_slide_ < *l*_bare_). To successfully skip, we here also need the roadblock footprint (*l*_foot_) to be smaller than length of the region not typically probed by sliding (*l*_foot_ < *l*_sNs_ − *l*_slide_). Taken together, the roadblock coverage (*ρ*_RB_) where sNs-search remains efficient is limited by

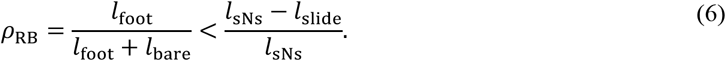

**Fig. 4.**
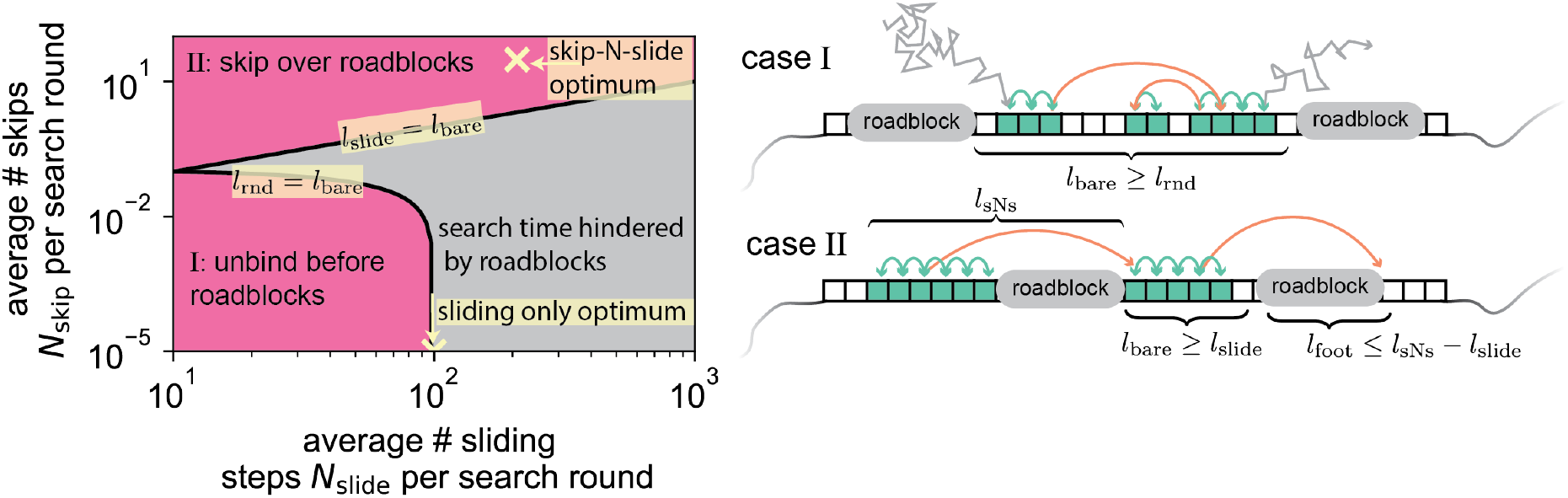
Effect of roadblocks on conditions for fast search. (left) The method for bypassing roadblocks indicated for varying *N*_slide_ and *N*_skip_. We have used *l*_skip_ = 30 nt and *τ*_3D_ = 10*τ*_skip_ = 100*τ*_slide_, and assume roadblocks are typically *l*_bare_ = 10 nt apart. Grey region indicates that roadblocks are so densely packed that they will force the protein to slide over sites already visited. In the top pink region (*l*_slide_ < *l*_bare_) the protein can use skips to move past roadblocks, and in the lower pink region in (*l*_rnd_ < *l*_bare_) proteins typically unbinds before encountering any roadblock. The location of local minima in search time (yellow markers) are the same as in **Fig. 3b.** (right) Illustration of the mechanism behind bypassing roadblocks in case I and case II.

For our measurements on Ago, the maximal coverage is roughly 70%. Typical mRNA 3’-UTR substrates are estimated to be 40-80% covered with proteins^35^, indicating that sNs-search could proceed largely unhindered.

## Discussion

We have presented both a general analytical theory and a single-molecule assay that quantitatively capture the distribution of skips taken during 1D lateral search at nucleotide resolution. By measuring the time it takes a searcher to transition between both close-by and far-apart traps on the substrate, we can always determine the ratio between sliding and skipping lengths (**Fig. 1**). Furthermore, if the transition time vs. trap separation has a local minimum, then the skips have a definite length that is given by the trap separation at this minimum. Applying the assay and theory to hAgo2 and CbAgo (**Fig. 2**), we found that both alternate between skipping about 30 nt and sliding 10 nt. As we find that short skips are suppressed, we argued that skips are likely realized by intersegmental transfer.

We further quantified the potential benefit of adopting an sNs-search strategy *in vivo* by first giving the general conditions under which the search time is minimized on bare substrates (**Fig. 3**). These conditions show that optimal sNs searchers always spend more time skipping and sliding along the substrate, than diffusing in 3D. Interestingly, for Ago there is only a narrow region of parameter space (intersection of the yellow slice and green region in **Fig. 3c**) where the speed of optimal sNs-search compares to that of optimal search without skips. Despite the small gain available on bare substrates, our measurements show that sNs-search does occur. We next established that optimal sNs-search can be sustained on crowded substrates up to a maximal coverage of roadblocks (**Equation 6**). Though there is no appreciable speedup for Ago using sNs-optimized search on bare substrates, it allows the protein to efficiently search along substrates with up to about 70% coverage (broadly consistent with the 40-80% coverage on typical 3’-UTR mRNA^35^).

Proteins searching along more rigid double-stranded DNA can similarly be investigated for skips. It would be particularly interesting to consider the *E.coli* lac repressor, as it has been shown to spend more than 50% of the time diffusing along the substrate^49,50^, which is consistent with an optimal sNs-search strategy. *In vivo*, skips may be especially important for search on chromatin, which adopts a variety of conformations, densities, and 3D motifs^51^. As cellular DNA and RNA are generally densely covered by roadblocks and adopt various 3D conformations, we expect many other target searchers to adopt sNs-search to rapidly find their target sites.

## Methods

### Protein purification

CbAgo was purified according to Hegge et al, 2019^45^. hAgo2 was purified according to Chandradoss et al, 2015^43^.

### Nucleic acid preparation

RNA constructs with a single amine-C6-uridine modification were ordered from STPharm. After labelling with Cy5 according to^52^ the constructs were precipitated. The RNA constructs were subsequently annealed to a DNA splint (specific for RNA and U40 mer), a second DNA splint (for ligating U40 mers) and a U40 mer (in the ratio 1:2:3:3). After ligation with T4 RNA ligase II (NEB), the ligated constructs were run on a 10% PAGE. Different ligated populations are created through this process (for example, TGT-U40 or TGT-U40-U40 etc.) and these are then excised from the gel and concentrated through ethanol precipitation. The concentrated and ligated RNA constructs were again annealed to a DNA construct and an RNA target with biotin on the 3’ end. Ligation was again performed with T4 RNA ligase II.

DNA oligos with a single amine-C6-thymine modification were ordered from ELLA Biotech GmbH and labeled in the same way as the RNA.

### Sample preparation

Quartz slides were prepared according to Chandradoss et al.^53^. Briefly, quartz slides were cleaned with detergent, sonicated and treated with acetone and subsequently KOH. Coverslips were directly sonicated with KOH. Piranha cleaning was done followed by treatment with methanol and incubation of (3-Aminopropyl)triethoxysilane (APTES) for both coverslips and quartz slides. PEGylation took place overnight and slides and coverslips were stored at −20°C. Before single-molecule experiments, an extra round of PEGylation took place with MSPEG-4.

The quartz slide was then assembled with scotch tape and epoxy glue and the chamber is flushed in T50 and 1% Tween-20 for >10min to further improve the surface quality of the single-molecule chambers^54^. Channels were thoroughly washed with T50 before adding in streptavidin (0.1 mg/mL) for 1 min. Subsequently, DNA or RNA was immobilized on the surface through biotin-streptavidin conjugation. 10 nM CbAgo or hAgo2 was incubated with 1 nM guide in (100 mM NaCl for CbAgo, 50 mM NaCl for hAgo2), 50 mM Tris, 1 mM Trolox, 0.8% glucose for ~30 min. Lastly, glucose oxidase (0.1 mg/mL final conc.) and catalase (17 μg/mL final conc.) were added and introduced in the chamber.

### Experimental setup

Single-molecule experiments were performed on a custom built inverted microscope (IX73, Olympus) using prism-TIRF and a 60X water immersion objective (UPLSAPO60XW, Olympus). The Cy3 dye was excited using a 532 nm diode laser Compass 215M/50mW, Coherent) and the Cy5 dye was excited using a 637 nm diode laser (OBIS 637 nm LX 140 mW). The scattered light was blocked by a 532 nm notch filter (NF03-532E-25, Semrock) and a 633 nm notch filter (NF03-633E-25, Semrock) after which the remaining signal from the fluophores was separated into two separate channels. Lastly, the light is projected on a EM-CCD camera (iXon Ultra, DU-897U-CS0-\# BV, Andor Technology).

Before each experiment, a reference movie was taken with the red laser to excite the Cy5 dyes on the nucleic acid molecules of interest. After that, a movie is taken with the green laser. The single-molecule experiments were taken at room temperature (20 ± 0.1 °C).

### Analysis of raw data

The raw data was analyzed using custom written code in IDL, where the reference movie is used to take into account only the regions of interest (i.e. the regions that contain a Cy5). The resulting time traces where further analyzed in MATLAB (Mathworks) where the shuttling rates were extracted through the use of Hidden Markov software called ebFRET (http://ebfret.github.io/) and custom written code in MATLAB.

## Acknowledgements

M.K. was supported by the Netherlands Organization for Scientific Research (NWO/OCW), as part of the Frontiers in Nanoscience program. M.D. acknowledges support from the Parents in KIND program, sponsored by The Kavli Institute of Nanoscience Delft, the Department of Bionanoscience at TU Delft, and through a Spinoza Prize awarded to M. Dogterom by NWO. C.J. was supported by Vidi (864.14.002) of the Netherlands Organization for Scientific research and an ERC Consolidator grant (819299) of the European Research Council. I.M. was supported by NIH grant R35GM127090.

## Author Contributions

M.K. and M.D. developed the analytical models. M.K. performed numerical simulations and fitting to data. T.J.C and C.J. designed the experiments. T.J.C. performed the measurements. I.M. provided purified hAgo2. All authors contributed to writing the manuscript.

## References

1. Berg, O. G., Winter, R. B. & von Hippel, P. H. Diffusion-driven mechanisms of protein translocation on nucleic acids. 1. Models and theory. Biochemistry 20, 6929–6948 (1981).

2. Elowitz, M. B. et al. Protein Mobility in the Cytoplasm of Escherichia coli. J. Bacteriol. 181, 197–203 (1999).

3. Halford, S. E. & Marko, J. F. How do site-specific DNA-binding proteins find their targets? Nucleic Acids Res. 32, 3040–3052 (2004).

4. Vonhippel, P. H. & Berg, O. G. Facilitated Target Location in Biological-Systems. J. Biol. Chem. 264, 675–678 (1989).

5. Riggs, A. D., Bourgeois, S. & Cohn, M. The lac represser-operator interaction. III. Kinetic studies. J. Mol. Biol. 53, 401–417 (1970).

6. Sheinman, M. & Kafri, Y. The effects of intersegmental transfers on target location by proteins. Phys. Biol. 016003, (2009).

7. Slutsky, M. & Mirny, L. A. Kinetics of Protein-DNA Interaction: Facilitated Target Location in Sequence-Dependent Potential. Biophys. J. 87, 4021–4035 (2004).

8. Hu, T. & Shklovskii, B. I. How a protein searches for its specific site on DNA: The role of intersegment transfer. Phys. Rev. E - Stat. Nonlinear, Soft Matter Phys. 76, 1–8 (2007).

9. Li, G., Berg, O. G. & Elf, J. Effects of macromolecular crowding and DNA looping on gene regulation kinetics. Nat. Phys. 5, 294–297 (2009).

10. Lomholt, M. A., van den Broek, B., Kalisch, S.-M. J., Wuite, G. J. L. & Metzler, R. Facilitated diffusion with DNA coiling. Proc. Natl. Acad. Sci. 106, 8204–8208 (2009).

11. Marcovitz, A. & Levy, Y. Obstacles May Facilitate and Direct DNA Search by Proteins. Biophys. J. 104, 2042–2050 (2013).

12. Krepel, D., Gomez, D., Klumpp, S. & Levy, Y. Mechanism of Facilitated Diffusion during a DNA Search in Crowded Environments. J. Phys. Chem. B 120, 11113–11122 (2016).

13. Vuzman, D. & Levy, Y. The “monkey-bar” mechanism for searching for the DNA target site: The molecular determinants. Isr. J. Chem. 54, 1374–1381 (2014).

14. Shvets, A. A., Kochugaeva, M. P. & Kolomeisky, A. B. Mechanisms of protein search for targets on DNA: Theoretical insights. Molecules 23, 1–18 (2018).

15. Shvets, A. A. & Kolomeisky, A. B. Crowding on DNA in Protein Search for Targets. J. Phys. Chem. Lett. 7, 2502–2506 (2016).

16. Bauer, M. & Metzler, R. Generalized facilitated diffusion model for DNA-binding proteins with search and recognition states. Biophys. J. 102, 2321–2330 (2012).

17. Blainey, P. C., van Oijen, A. M., Banerjee, A., Verdine, G. L. & Xie, X. S. A base-excision DNA-repair protein finds intrahelical lesion bases by fast sliding in contact with DNA. Proc. Natl. Acad. Sci. 103, 5752–5757 (2006).

18. Bonnet, I. et al. Sliding and jumping of single EcoRV restriction enzymes on non-cognate DNA. Nucleic Acids Res. 36, 4118–4127 (2008).

19. Ragunathan, K., Liu, C. & Ha, T. RecA filament sliding on DNA facilitates homology search. Elife 1–14 (2012). doi:10.7554/eLife.00067

20. Stanford, N. P., Szczelkun, M. D., Marko, J. F. & Halford, S. E. One- and three-dimensional pathways for proteins to reach specific DNA sites. EMBO J. 19, 6546–6557 (2000).

21. Tafvizi, A., Huang, F., Fersht, A. R., Mirny, L. A. & van Oijen, A. M. A single-molecule characterization of p53 search on DNA. Proc. Natl. Acad. Sci. 108, 563–568 (2011).

22. Wang, Y. M., Austin, R. H. & Cox, E. C. Single molecule measurements of repressor protein 1D diffusion on DNA. Phys. Rev. Lett. 97, 1–4 (2006).

23. Zandarashvili, L. et al. Balancing between affinity and speed in target DNA search by zinc-finger proteins via modulation of dynamic conformational ensemble. Proc. Natl. Acad. Sci. 112, E5142–E5149 (2015).

24. Gowers, D. M., Wilson, G. G. & Halford, S. E. Measurement of the contributions of 1D and 3D pathways to the translocation of a protein along DNA. Proc. Natl. Acad. Sci. 102, 15883–15888 (2005).

25. Graneli, A., Yeykal, C. C., Robertson, R. B. & Greene, E. C. Long-distance lateral diffusion of human Rad51 on double-stranded DNA. Proc. Natl. Acad. Sci. 103, 1221–1226 (2006).

26. Hammar, P. et al. The lac Repressor Displays Facilitated. Science (80-.). 336, 1595–1598 (2012).

27. Kim, J. H. & Larson, R. G. Single-molecule analysis of 1D diffusion and transcription elongation of T7 RNA polymerase along individual stretched DNA molecules. Nucleic Acids Res. 35, 3848–3858 (2007).

28. Kochaniak, A. B. et al. Proliferating cell nuclear antigen uses two distinct modes to move along DNA. J. Biol. Chem. 284, 17700–17710 (2009).

29. Koh, H. R., Kidwell, M. A., Doudna, J. & Myong, S. RNA scanning of a molecular machine with a built-in ruler. J. Am. Chem. Soc. 139, 262–268 (2017).

30. Leith, J. S. et al. Sequence-dependent sliding kinetics of p53. Proc. Natl. Acad. Sci. 109, 16552–16557 (2012).

31. Normanno, D. et al. Probing the target search of DNA-binding proteins in mammalian cells using TetR as model searcher. Nat. Commun. 6, (2015).

32. Cui, T. J. et al. Argonaute bypasses cellular obstacles without hindrance during target search. Nat. Commun. 10, 4390 (2019).

33. Brogaard, K., Xi, L., Wang, J. P. & Widom, J. A map of nucleosome positions in yeast at base-pair resolution. Nature 486, 496–501 (2012).

34. Azam, T. A. L. I., Iwata, A., Nishimura, A., Ueda, S. & Ishihama, A. Growth Phase-Dependent Variation in Protein Composition of the. Society 181, 6361–6370 (1999).

35. Silverman, I. M. et al. RNase-mediated protein footprint sequencing reveals protein-binding sites throughout the human transcriptome. Genome Biol. 15, 1–16 (2014).

36. van den Broek, B., Lomholt, M. A., Kalisch, S.-M. J., Metzler, R. & Wuite, G. J. L. How DNA coiling enhances target localization by proteins. Proc. Natl. Acad. Sci. 105, 15738–15742 (2008).

37. Bartel, D. P. Metazoan MicroRNAs. Cell 173, 20–51 (2018).

38. Hutvagner, G. & Simard, M. J. Argonaute proteins: Key players in RNA silencing. Nat. Rev. Mol. Cell Biol. 9, 22–32 (2008).

39. Gebert, L. F. R. & MacRae, I. J. Regulation of microRNA function in animals. Nat. Rev. Mol. Cell Biol. 20, 21–37 (2019).

40. Hegge, J. W., Swarts, D. C. & Van Der Oost, J. Prokaryotic argonaute proteins: Novel genome-editing tools? Nat. Rev. Microbiol. 16, 5–11 (2018).

41. Swarts, D. C. et al. The evolutionary journey of Argonaute proteins. Nat. Struct. Mol. Biol. 21, 743–753 (2014).

42. Sternberg, S. H., Redding, S., Jinek, M., Greene, E. C. & Doudna, J. A. DNA interrogation by the CRISPR RNA-guided endonuclease Cas9. Nature 507, 62–67 (2014).

43. Chandradoss, S. D., Schirle, N. T., Szczepaniak, M., Macrae, I. J. & Joo, C. A Dynamic Search Process Underlies MicroRNA Targeting. Cell 162, 96–107 (2015).

44. Globyte, V., Lee, S. H., Bae, T., Kim, J. & Joo, C. CRISPR/Cas9 searches for a protospacer adjacent motif by lateral diffusion. EMBO J. 38, e99466 (2019).

45. Hegge, J. W. et al. DNA-guided DNA cleavage at moderate temperatures by Clostridium butyricum Argonaute. Nucleic Acids Res. 47, 5809–5821 (2019).

46. Schirle, N. T., Sheu-Gruttadauria, J. & Macrae, I. J. Structural basis for microRNA targeting. Science (80-.). 346, (2014).

47. Klein, M., Chandradoss, S. D., Depken, M. & Joo, C. Why Argonaute is needed to make microRNA target search fast and reliable. Semin. Cell Dev. Biol. 65, 20–28 (2017).

48. Cuculis, L., Abil, Z., Zhao, H. & Schroeder, C. M. Direct observation of TALE protein dynamics reveals a two-state search mechanism. Nat. Commun. 6, 1–11 (2015).

49. Elf, J., Li, G.-W. & Xie, X. S. Probing Transcription Factor Dynamics at the Single-Molecule Level in a Living Cell. Science (80-.). 316, 1191–1195 (2007).

50. Kao-Huang, Y. et al. Nonspecific DNA binding of genome-regulating proteins as a biological control mechanism: Measurement of DNA-bound Escherichia coli lac repressor in vivo. Proc. Natl. Acad. Sci. 74, 4228–4232 (1977).

51. Ou, H. D. et al. ChromEMT: Visualizing 3D chromatin structure and compaction in interphase and mitotic cells. Science (80-.). 357, (2017).

52. Joo, C. & Ha, T. Single-molecule FRET with total internal reflection microscopy. Cold Spring Harb. Protoc. 7, 1223–1237 (2012).

53. Chandradoss, S. D. et al. Surface Passivation for Single-molecule Protein Studies. J. Vis. Exp. 1–8 (2014). doi:10.3791/50549

54. Pan, H., Xia, Y., Qin, M., Cao, Y. & Wang, W. A simple procedure to improve the surface passivation for single molecule fluorescence studies. Phys. Biol. 12, (2015).

