## Supplementary Information for "Skipping and sliding to optimize target search on protein-bound DNA and RNA"

### 1 Modeling sNs lateral diffusion

To explain the experimental data shown in **Figs.** 2c and d, we build a kinetic model for the lateral diffusion by a target searching protein that both skips and slides. We imagine our searcher being capable of binding at any position, and when bound to site $i$, the it diffuses away (in either direction) at a rate

$$k_{\text{move},i} = \begin{cases} k_{\text{trap}} & \text{at trap} \\ k_{\text{ns}} & \text{at non-specific site.} \end{cases} \tag{S1}$$

Based on the fact that we cannot resolve actual diffusion in experiments (**Fig.** 2b) we assume that sliding between non-specific sites is near instantaneous, and that we only need to keep track of the total time spent in the initial trap before being captured by the final trap. Furthermore, as we typically see more than 10 shuttle events occur prior to unbinding, we will ignore the possibility of unbinding within one shuttling event. In every move, the protein can either slide to one of its neighbors, or skip some bases and land further away. We assign a probability that such a step is of (directed) length $l$ (in nucleotides)

$$S_l = \frac{l_{\text{slide}}^2}{1 + l_{\text{slide}}^2}\frac{1}{2}\delta_{|l|,1} + \frac{1}{1 + l_{\text{slide}}^2}s_l. \tag{S2}$$

Here $\delta$ denotes the Kronecker delta and $s_l$ represents the (normalized) symmetric distribution of skip distances (see orange peaks in **Fig.** 1b). The weight of each term in **Equation** S2 are such that we on average have $l_{\text{slide}}^2$ sliding steps between each skip.

As we show below, the shuttling time at small and large trap separations turns out to be independent of many of the details of the skip distribution $s_l$. To be able to capture the essence of all the possible skipping methods, numerical results are obtained assuming $s_l$ to be a bi-modal Gaussian mixture model, with equal-weight peaks at $\pm\mu_{\text{skip}}$, both with variance $\sigma_{\text{skip}}^2$.

#### 1.1 Asymptotic theory at large trap separations

In a previous paper we captured the shuttling time in the limit of small trap separations [1], and here we turn to the limit of large trap separation. To analytically capture the behavior of the shuttling time for large trap separations, we consider the limit where there are typically many sNs cycles ($l_{\text{slide}}^2$ sliding steps and a skip) in each shuttling event. As we do not account for the time spent sliding and skipping (sub resolution), the time to shuttle should be equal to

$$T_{\text{shuttle}}(d_{\text{trap}}) = N_{\text{trap}}\tau_{\text{trap}}. \tag{S3}$$

Here $d_{\text{trap}}$ is the trap distance, $\tau_{\text{trap}}$ is the average trap escap time, and $N_{\text{trap}}$ is the average number of retrapping events before getting trapped in the second trap; with the latter being a large number for large trap separations.

We move to a coarse grained system, in which each sNs cycle moves a ms distance $l_{\text{sNs}}^2 = l_{\text{slide}}^2 + \mu_{\text{skip}}^2 + \sigma_{\text{skip}}^2$, and interrogates the fraction $\rho_{\text{sNs}} = l_{\text{slide}}/l_{\text{sNs}}$ of the bases it passes over. After the first skip, the protein has moved to a substrate segment that lies $l_{\text{sNs}}$ away from the initial trap. To get retrapped, the system has to return to the segment containing the first trap, and then visit the trap. We have previously shown that when using sliding alone, the probability to shuttle (before getting recaptured) once the initial trap is escaped, equals the inverse of the number of steps between the traps [1]. We can lift this result to the coarse grained description by noting that the probability to shuttle between trap segments rather than returning to the initial trap segment is $P_{\text{shuttle}} = l_{\text{sNs}}/d_{\text{trap}}$.

After jumping out of the first segment with the first skip (shown in yellow in **Supplementary Figure** 2a), the probability return to the initial segment ($P_{\text{return}}$) without being trapped by the second trap can be calculated by summing probabilities over all possible paths that realize this

$$\begin{aligned} P_{\text{segment}} &= (1 - P_{\text{shuttle}}) \\ &+ P_{\text{shuttle}} \sum_{m=0}^{\infty} \left( (1 - \rho_{\text{sNs}}) (1 - P_{\text{shuttle}}) \right)^m (1 - \rho_{\text{sNs}}) P_{\text{shuttle}} \\ &= 1 - \frac{P_{\text{shuttle}} \rho_{\text{sNs}}}{P_{\text{shuttle}} + \rho_{\text{sNs}} - P_{\text{shuttle}} \rho_{\text{sNs}}}. \end{aligned} \tag{S4}$$

The first term after the first equality corresponds to returning to the segment with the first trap directly (path a in **Supplementary Figure** 2a), and the other terms corresponds to shuttling over to the segment with the second trap in it (first factor, path b in **Supplementary Figure** 2a), but no get caught in the trap (second factor, path c in **Supplementary Figure** 2a), even after cycling back and forth between the trap and the segment just left of it another $m$ times (central $m$ factors, repetition of paths c and d in **Supplementary Figure** 2a), before shuttling back straight to the segment with the initial trap (last factor, path e in b in **Supplementary Figure** 2a). Given the probability of re-entering the first segment, the average number of times this occurs prior to eventually shuttling across equals

$$\begin{aligned} n_{\text{segment}} &= \sum_{n=0}^{\infty} n P_{\text{segment}}^n (1 - P_{\text{segment}}) \\ &= \frac{P_{\text{segment}}}{1 - P_{\text{segment}}} = \frac{d_{\text{trap}}}{l_{\text{sNs}}} + \frac{l_{\text{sNs}}}{l_{\text{slide}}} - 2. \end{aligned} \tag{S5}$$

We next estimate the average number of times $n_{\text{recapture}}$ the protein gets recaptured back to the original trap once it has come back to the original segment. Assuming that a sufficient number of sNs cycles have taken place, the protein's position is uniformly spread throughout the $l_{\text{sNs}}$ long segment upon landing back in it. Given there are typically $l_{\text{slide}}^2$ steps taken prior to skipping out, each with probability $1/l_{\text{sNs}}$ to be landing the system in the trap, we on average have $n_{\text{recapture}} = l_{\text{slide}}^2 / l_{\text{sNs}}$ recapture events when returning to the first segment. The estimate of recapturing times will not be correct for the first few returns to the initial segment, since the return position might then be strongly correlated with the trap position.The time to de-correlate is not dependent on the trap separation though, so this can simply be corrected for with a constant term in the total number of re-trapping events

$$N_{\text{trap}} = \text{const.} + n_{\text{segment}} n_{\text{recapture}} = \text{const.} + \rho_{\text{sNs}}^2 d_{\text{trap}}. \tag{S6}$$

Hence, when placed sufficiently far apart, the shuttling time (**Equation** S3) should grow linearly with trap separation according to

$$T_{\text{shuttle}}(d_{\text{trap}}) = \text{const.} + \rho_{\text{sNs}}^2 \tau_{\text{trap}} d_{\text{trap}}. \tag{S7}$$

To validate our scaling argument we next solve the stochastic hopping model numerically.

##### 1.1.1 Numerically solving for shuttling time

Every shuttling event starts at $t = 0$ with the protein bound at one of the two traps, and ends at time $t = T_{\text{shuttle}}$ when the searcher for the first time reaches the other trap, located $d_{\text{trap}}$ sites away. For ease of notation we assume no flanks to either trap, but note that the results can be trivially extended to the case with flanks by doubling the effective trapping time to account for that half the trap escapes go in the wrong direction, and will thus always be re-trapped before shuttling. Letting $P_i(t)$ denote the probability for the protein to reside at site $i$ at time $t$, we define the state vector

$$\vec{P}(t) = \left[ P_1(t), P_2(t), ..., P_{d_{\text{trap}}-1}(t) \right]^T. \tag{S8}$$

Here we include the probabilities of all but the second trapped state, as this is simply set by normalization. To capture the first-passage time [2] into the second trap, we artificially make the last trap absorbing, and we can write the Master Equation controlling the dynamics as.

$$\frac{\partial \vec{P}(t)}{\partial t} = -K \vec{P}(t) \quad \Rightarrow \quad \vec{P}(t) = e^{-Kt} \vec{P}(0) \tag{S9}$$

where we have defined the rate matrix $K$ through

$$K_{j,i} = \begin{cases} -k_{\text{move},i} S_{i-j}, & 1 \le i \ne j \le d_{\text{trap}} - 1 \\ \sum_{l=1,\, l\neq i}^{d_{\text{trap}}-1} K_{l,i}, & 1 \le i = j \le d_{\text{trap}} - 1. \end{cases} \tag{S10}$$

The shuttle event starts with the protein located at the first trap, $P_i(0) = \delta_{i0}$, and ends when the second trap is reached. The first passage time into the second trap then equals the average time to reach the absorbing state. Noting that what ever ends up in the absorbing state in the time interval $[t, t+dt]$ also transitioned to this state for the first time, we can write the first-passage time probability distribution from one trap to another as

$$\Psi_{\text{shuttle}}(\tau) = \frac{\partial P_{d_{\text{trap}}}(\tau)}{\partial \tau} = -\sum_{j=1}^{d_{\text{trap}}-1} \frac{\partial P_j(\tau)}{\partial \tau}. \tag{S11}$$

The average shuttling time can now be written as

$$\begin{aligned} T_{\text{shuttle}} &= \int_0^\infty \tau \Psi_{\text{shuttle}}(\tau) \mathrm{d}\tau = -\sum_{j\neq d_{\text{trap}}} \int_0^\infty \tau \frac{\partial P_j(\tau)}{\partial \tau} \mathrm{d}\tau \\ &= \sum_{j\neq d_{\text{trap}}} \int_0^\infty P_j(\tau) \mathrm{d}\tau = \sum_{j\neq d_{\text{trap}}} \left( \int_0^\infty e^{-K\tau} \mathrm{d}\tau \right) P_j(0) = \sum_{j\neq d_{\text{trap}}} (K^{-1} \vec{P}(0))_j. \end{aligned} \tag{S12}$$

From this we see that we can calculate the average shuttling time through numerical inversion of our rate matrix. To validate our scaling arguments for large trap separations we now calculate the final slope of $T_{\text{shuttle}}(d_{\text{trap}})$ using **Equation** S12 for a wide range of $l_{\text{slide}}$, $\mu_{\text{skip}}$ and $\sigma_{\text{skip}}$. In **Supplementary Figure** 2b we show our numerical results for $l_{\text{slide}} \in$ [1 nt, 6 nt, 12 nt, 18 nt, 24 nt, 30 nt, 36 nt, 42 nt], $\mu_{\text{skip}} \in$ [0 nt,6 nt, 12 nt, 18 nt, 24 nt, 30 nt, 36 nt, 42 nt] and $\sigma_{\text{skip}} \in$ [0.01 nt, 6 nt, 12 nt, 18 nt, 24 nt, 30 nt, 36 nt, 42 nt]. The distance between traps varied from 1-250 nt. As can be seen from **Supplementary Figure** 2b we have complete data collapse for the parameter range considered.

##### 1.2 Bootstrapping for error estimation and based on smFRET data

Fitting the data from the tandem target assay to **Equation** 1 provides the estimate of $\tau_{\text{trap}}$. We bootstrapped the dwell time distributions acquired using the original tandem target assay (distances of 11 nt, 15 nt, 18 nt and 22 nt (CbAgo) and 7 nt, 11 nt, and 15 nt (hAgo2)). For each of the $10^5$ bootstrap samples we calculated new values for the associated $T_{\text{shuttle}}$'s and repeated the fit to **Equation** 1 to obtain an error estimate in the fitted value of the escape rate. In similar fashion, we used **Equation** S7, together with the estimate of $\tau_{\text{trap}}$ from the original dataset, to determine $\rho_{\text{sNs}}$ (distances of 64 nt, 92 nt and 120 nt (CbAgo) and 80 nt, 120 nt, 160 nt (hAgo2)). All analysis was performed with a custom code written in Python. Shaded areas in **Figs.** 2c and d represent 95% confidence intervals.

#### 2 Optimal search times for sNs lateral diffusion

Here we determine the time needed for a protein to locate a single target embedded among $L$ non-specific binding sites. We seek to understand how the balance between skipping allows the searcher to reach distant targets without causing the searcher to miss the target. In one 1D search round, we take the protein to bind and undergo an average of $N_{\text{skip}}$ sNs-cycles before dissociating, meaning a total of $N_{\text{slide}} = N_{\text{skip}} l_{\text{slide}}^2$ sliding steps have been performed. The average time needed for each search round $T_{\text{rnd}} = \tau_{1\text{D}} + \tau_{3\text{D}}$ can be split into the time $\tau_{1\text{D}}$ spent interrogating sequences through 1D lateral diffusion and the time $\tau_{3\text{D}}$ to return from solution to interrogate a base. The 1D lateral diffusion time $\tau_{1\text{D}} = T_{\text{slide}} + T_{\text{skip}}$ can be further split into the total time spent sliding and interrogating off-targets $T_{\text{slide}} = N_{\text{slide}} \tau_{\text{slide}}$ and the total time spent skipping and interrogating the landing sites $T_{\text{skip}} = N_{\text{skip}} \tau_{\text{skip}}$.

The average number of sites checked in one 1D lateral diffusion event is $l_{\rm rnd}\rho_{\rm rnd}$, and we need an average number $N_{\rm rnd} = L/(l_{\rm rnd}\rho_{\rm rnd})$ of rounds to find one target among $L$ sites.

$$T_{\rm search} = T_{\rm rnd}N_{\rm rnd} = \left( \overbrace{\underbrace{N_{\rm slide}\tau_{\rm slide}}_{T_{\rm slide}} + \underbrace{N_{\rm skip}\tau_{\rm skip}}_{T_{\rm skip}}}^{\tau_{\rm 1D}} + \tau_{\rm 3D} \right) \frac{L}{l_{\rm rnd}\rho_{\rm rnd}} \tag{S13}$$

Note that lowering any of the microscopic timescales associated with the different search modes will always speed up the search. We therefore take these microscopic times to already be reduced as far as allowed by external constraints. For skipping to ever be beneficial over 3D diffusion $\tau_{\rm 3D} > \tau_{\rm skip}$, and for sliding to ever be beneficial over skipping we need $\tau_{\rm skip} > \tau_{\rm slide}$. We here assume $\tau_{\rm 3D} > \tau_{\rm skip} > \tau_{\rm slide}$ in order to focus on the non-trivial case. Similarly, increasing only the mean-squared skipping distance $l_{\rm skip}^2$ will always reduce the scanning redundancy, and so will always reduce the search time. Since we observe skips of finite length, we will also assume the skipping length to be externally limited, and take it to be fixed. In the end, we are left with two adjustable parameters that could have been further optimized by evolution: the number of skips $N_{\rm skip}$ and the total number $N_{\rm slide} = N_{\rm skip}l_{\rm slide}^2$ of sliding steps in one search round.

#### 2.1 The effective scanning density $\rho_{\rm rnd}$

Before optimizing **Equation** S13 we need to determine the effective scanning density $\rho_{\rm rnd}$. We find an approximate form for the effective scanning density by considering the shuttling to a segment $l_{\rm rnd}$ away as a proxy for free sNs diffusive motion between binding and unbinding. We use Monte Carlo simulations to show that the approximation remains valid over the relevant parameter ranges (**Supplementary Figure** 4).

Assume the protein starts its sNs search in the first segment and ends it the first time it reaches $D = l_{\rm rnd}/l_{\rm sNs}$ (**Supplementary Figure** 4a). To calculate the average probability to interrogate any site along the way, we first determine the probability $p_{\rm no\ check}$ of missing a site located within an intermediate segment $D_{\rm check} < D$, prior to reaching $D$. During an sNs cycle, each site will get visited/interrogated with probability $\rho_{\rm sNs}$. Hence, to skip past a site within $D_{\rm check}$, the system first reaches segment $D_{\rm check}$ (path a in **Supplementary Figure** 4a), and can then escape without interrogation (path b in **Supplementary Figure** 4a) and return (path c in **Supplementary Figure** 4a) any number of times, before finally skipping away and make its way to segment $D$ (path d in **Supplementary Figure** 4a). Summing over these paths, we have the probability that a specific base in segment $D_{\rm check}$ is checked as

$$\begin{aligned} p_{\rm no\ check}(D_{\rm check}) &= 1 \times \sum_{m=0}^{\infty} \left[\frac{1}{2}(1-\rho_{\rm sNs}) \times 1 + \frac{1}{2}(1-\rho_{\rm sNs}) \times (1 - P_{\rm shuttle})\right]^m \frac{1}{2}(1-\rho_{\rm sNs})P_{\rm shuttle} \\ &= \frac{(1-\rho_{\rm sNs})/(2\rho_{\rm sNs})}{D - D_{\rm check} + (1-\rho_{\rm sNs})/(2\rho_{\rm sNs})}. \end{aligned} \tag{S14}$$

Here we have used that $P_{\rm shuttle} = 1/(D - D_{\rm check})$ corresponds to the shuttling from segment $D_{\rm check} + 1$ to segment $D = l_{\rm rnd}/l_{\rm sNs}$. The probability to interrogate a particular site within $D_{\rm check}$ at least once prior to reaching segment $D$ equals

$$\begin{aligned} p_{\rm check}(D_{\rm check}) &= 1 - p_{\rm no\ check}(D_{\rm check}) \\ &= \frac{D - D_{\rm check}}{D - D_{\rm check} + (1-\rho_{\rm sNs})/(2\rho_{\rm sNs})}. \end{aligned} \tag{S15}$$

The average probability to interrogate any particular site between sites 1 and $l_{\rm rnd}$ at least once prior to unbinding, $\rho_{\rm rnd}$, is obtained after further approximating the sum over $D_{\rm check}$ positions as an integral

$$\rho_{\rm rnd} \approx \frac{1}{D}\int_0^D p_{\rm check}(D_{\rm check})\mathrm{d}D_{\rm check} = 1 - \frac{\ln(1+2x)}{2x} \tag{S16}$$

with

$$x = \frac{\rho_{\rm sNs}}{1-\rho_{\rm sNs}}\sqrt{N_{\rm skip}} = \frac{\rho_{\rm sNs}}{1-\rho_{\rm sNs}}\sqrt{l_{\rm rnd}/l_{\rm sNs}} \tag{S17}$$

To validate our approximation of the effective scanning density, we next turn to Montecarlo simulations.

##### 2.1.1 Monte Carlo simulations for validating the effective scanning density

To test the validity of **Equation** S16, we set up Monte Carlo simulations (code written in Python). After landing on our sNs coarse grained lattice, the proteins are assigned a unity step rate to either side (i.e. we her work in the units of the sliding rate), as well as an unbinding rate $u$. In every move, the protein diffuses to one of its neighboring site with a probability $\frac{1}{2+u}$ and unbinds with a probability $\frac{u}{2+u}$, meaning it performs on average $N_{\rm skip} = 2/u + 1$ skips before unbinding. When landing at a site, the protein interrogates it to find the target with a fixed probability $\rho_{\rm sNs}$. Each of the 1000 runs ends when the protein stochastically unbinds.

The achieve data collapse according to **Equation** S16, we need to calculate the $\rho_{\rm rnd}$ and match it to $x$ in accordance with **Equation** S17. We estimate the value of $\rho_{\rm rnd}$ as the fraction of sites visited that are also interrogated, and measure the total excursion length $l_{\rm rnd}$ for each run to calculate $x$. Error bars in **Supplementary Figure** 4b show 95% confidence intervals for both $x$ and $\rho_{\rm sNs}$. Simulations were repeated for $\rho_{\rm sNs}$ in [$10^{-10}$, $10^{-9}$,$10^{-8}$,...,$10^{-2}$, 0.9,0.8,...0.1], and $u$ in [$10^{-5}$, $10^{-4}$, $10^{-3}$, $10^{-2}$] (or equivalently $N_{\rm skip} \approx [2\cdot 10^5, 2\cdot 10^4, 2\cdot 10^3, 2\cdot 10^2]$). **Supplementary Figure** 4b compares **Equation** S16 (solid line) to Monte Carlo simulations (markers) in which a fixed average number of skips is set, and the complete data collapse shows that our approximation was justified.

#### 2.2 Optimizing the search time

For ease of calculation, we change the independent variables to

$$x = \frac{\rho_{\rm sNs}}{1-\rho_{\rm sNs}}\sqrt{N_{\rm skip}} \quad , \quad y = \frac{\rho_{\rm sNs}}{1-\rho_{\rm sNs}} \tag{S18}$$

and minimize the search time (**Equation** S13) with respect to these. In terms of these variables, the total times spent on sliding and skipping becomes

$$T_{\rm slide} = N_{\rm slide}\tau_{\rm slide}, \quad N_{\rm slide} = \frac{(x l_{\rm skip})^2}{1+2y} \tag{S19}$$

$$T_{\rm skip} = N_{\rm skip}\tau_{\rm skip}, \quad N_{\rm skip} = \left(\frac{x}{y}\right)^2. \tag{S20}$$

As the root-mean-squared excursion during a search round can be written as,

$$l_{\rm rnd} = \sqrt{N_{\rm slide} + l_{\rm skip}^2 N_{\rm skip}} = \sqrt{N_{\rm skip}}\, l_{\rm sNs} = \frac{(y+1)l_{\rm skip}}{y\sqrt{1+2y}}, \tag{S21}$$

and the number of rounds needed before finding the target site equals $N_{\rm rnd} = {}^{L}/_{l_{\rm rnd}\rho_{\rm rnd}(x)}$, the search time can be written as

$$\begin{aligned} T_{\rm search} &= L\frac{N_{\rm skip}\tau_{\rm skip} + N_{\rm slide}\tau_{\rm slide} + \tau_{\rm 3D}}{l_{\rm rnd}\rho_{\rm rnd}(x)} \\ &= L\frac{(xy l_{\rm skip})^2\tau_{\rm slide} + (1+2y)(x^2\tau_{\rm skip} + y^2\tau_{\rm 3D})}{\sqrt{1+2y}(1+y)yx\rho_{\rm rnd}(x) l_{\rm skip}}. \end{aligned} \tag{S22}$$

We start by examining the boundary of no skips, and then move on to the case with finite skips.

##### 2.2.1 The sliding optimum and its conditions

One minimum $T_{\text{search}}$ is found at the boundary $N_{\text{skip}} \to 0$, for which $x \to {}^1/_{N_{\text{skip}}} \to \infty$ (**Equation** S18), and $\rho_{\text{rnd}} \to 1$ (**Equation** S16), and $l_{\text{rnd}} \to \sqrt{N_{\text{slide}}}$ (**Equation** S21). This applied to **Equation** S22 results in the search time

$$T_{\text{search}} \to L\frac{N_{\text{slide}}\tau_{\text{slide}} + \tau_{3\text{D}}}{\sqrt{N_{\text{slide}}}} \tag{S23}$$

This can be optimized with respect to $N_{\text{slide}}$, resulting in the known condition [**reference Slutsky&Mirny, etc.**]

$$\tau_{3\text{D}} = \tau_{\text{slide}}N_{\text{slide}}^{\text{sliding}} = \tau_{1\text{D}} \quad \Rightarrow \quad N_{\text{slide}}^{\text{sliding}} = \frac{\tau_{3\text{D}}}{\tau_{\text{slide}}} \tag{S24}$$

where $N_{\text{slide}}^{\text{sliding}}$ is the optimal number of sliding steps before unbinding when only sliding is allowed. Calculating the derivative of the total search time with respect to $N_{\text{skip}}$ at constant $N_{\text{slide}}$

$$\left.\frac{\partial T_{\text{search}}}{\partial N_{\text{skip}}}\right|_{\substack{N_{\text{slide}}=N_{\text{slide}}^{\text{sliding}}\\ N_{\text{skip}}=0}} = \frac{1}{2}\frac{\tau_{\text{skip}} - \tau_{\text{slide}}}{\tau_{3\text{D}}}T_{\text{search}} \tag{S25}$$

show that this optimum is a local minimum as long as $\tau_{\text{skip}} > \tau_{\text{slide}}$, which we have already argued is the region we are interested in. At this sliding minimum, the total search time **Equation** S23 is

$$T_{\text{search}}^{\text{sliding}} = 2L\sqrt{\tau_{\text{slide}}\tau_{3\text{D}}}. \tag{S26}$$

##### 2.2.2 The sNs optimum and its conditions

When including skipping, we seek an optimal search time within the interior of the $\{x, y\}$-domain. Differentiating the search time (**Equation** S22) with respect to $x$ and $y$, and enforcing the zero condition on the derivatives we arrive at the general conditions for any optimum

$$\frac{\tau_{1\text{D}}}{T_{\text{rnd}}} = \frac{1}{2}\left(1 + x\partial_x \log\rho_{\text{rnd}}(x)\right) \tag{S27}$$

$$\frac{T_{\text{skip}}}{T_{\text{rnd}}} = \frac{1}{2}\frac{1}{1+y} + \frac{y}{1+2y}\left(\frac{1}{2} - \frac{T_{\text{slide}}}{T_{\text{rnd}}}\right) \tag{S28}$$

These are conditions that must be satesfied by any optimal strategy with finite skips, and since $\rho_{\text{rnd}}(x)$ increases with $x$ (**Equation** S16), we see directly from **Equation** S27 that any optimized searcher must spend more of its time skipping and sliding than diffusing in 3D.

##### 2.2.3 The sNs optimum in the low scanning-density regime

In order for skipping to be beneficial, the extra time needed to skip must be compensated by a decrease in redundancy. Therefore, a skip needs to bring the searcher away from the region just scanned by sliding. To elucidate this scenario, we consider the low scanning density limit where $l_{\text{skip}} \gg l_{\text{slide}}$, or equivalently $\rho_{\text{sNs}} \ll 1$ or $y \ll 1$. In this limit, **Equation** S27and S28 directly yields

$$\frac{T_{\text{slide}}}{T_{\text{rnd}}} = \frac{1}{2}x\partial_x \log\rho_{\text{rnd}}(x) \tag{S29}$$

$$\frac{T_{\text{skip}}}{T_{\text{rnd}}} = \frac{1}{2}. \tag{S30}$$

The above shows that at any local optimum in the low scanning density regime, half of the time is spent skipping. This should be compared to the high scanning-density regime containing the sliding optimum, where half of the time is spent sliding. Using **Equations** S36 and S20 to rewrite **Equation** S30 we have the following condition on the location of the optimum

$$(y^{\rm sNs}/y_0)^2 = \frac{(x^{\rm sNs}/x_0)^2}{1+(x^{\rm sNs}/x_0)^2}, \tag{S31}$$

$$x\partial_x \log \rho_{\rm rnd}(x^{\rm sNs}) = \frac{(x^{\rm sNs}/x_0)^2}{1+(x^{\rm sNs}/x_0)^2} \tag{S32}$$

where we have introduced

$$y_0 = \sqrt{\frac{\tau_{\rm skip}}{l^2_{\rm skip}\tau_{\rm slide}}}, \quad x_0 = \sqrt{\frac{\tau_{\rm 3D}}{l^2_{\rm skip}\tau_{\rm slide}}}. \tag{S33}$$

and used the fact that $T_{\rm skip} \to (xl_{\rm skip})^2\tau_{\rm slide}$ for small $y$. **Equation** S32 depends only on $x^{\rm sNs}$, and can be solved first for $x^{\rm sNs}$, and then **Equation** S31 directly gives $y^{\rm sNs}$. Next we solve **Equation** S32 in the limits of large and small $x^{\rm sNs}$.

$$x^{\rm sNs} = \begin{cases} (3/4)^{1/3}x_0^{2/3}, & x_0 \lll 1 \\ (\frac{\ln(x0)}{3})^{1/3}x_0^{2/3}, & 1 \ll x_0 \end{cases} \tag{S34}$$

Up to small numerical and logaritmic corrections we can therfore take

$$x^{\rm sNs} \approx x_0^{2/3} = \left(\frac{\tau_{\rm 3D}}{l^2_{\rm skip}\tau_{\rm slide}}\right)^{1/3}, \quad y^{\rm sNs} \approx \frac{y_0}{\sqrt{1+x_0^{2/3}}} = \sqrt{\frac{\frac{\tau_{\rm skip}}{l^2_{\rm skip}\tau_{\rm slide}}}{1+\left(\frac{\tau_{\rm 3D}}{l^2_{\rm skip}\tau_{\rm slide}}\right)^{1/3}}} \tag{S35}$$

We now have an expression for the optimal sNs system, and using this in **Equations** S36 and S20 yields e.g. the partition between skips and slides as

$$\frac{N^{\rm sNs}_{\rm skip}}{N^{\rm sNs}_{\rm slide}} = \frac{\tau_{\rm slide}}{\tau_{\rm skip}}\left(1+\left(\frac{\tau_{\rm 3D}}{l^2_{\rm skip}\tau_{\rm slide}}\right)^{1/3}\right) \tag{S36}$$

as well as the total search time at the sNs optimum in the low scanning density regime

$$T^{\rm sNs}_{\rm search} = 2L\sqrt{\tau_{\rm skip}\tau_{\rm slide}}\frac{x_0^{2/3}\sqrt{1+x_0^{2/3}}}{\rho_{\rm rnd}(x_0^{2/3})} = \begin{cases} 2L\sqrt{\tau_{\rm skip}\tau_{\rm slide}}, & \tau_{\rm 3D} \ll l^2_{\rm skip}\tau_{\rm slide} \\ \frac{2L}{l_{\rm skip}}\sqrt{\tau_{\rm skip}\tau_{\rm 3D}}, & \tau_{\rm 3D} \gg l^2_{\rm skip}\tau_{\rm slide}. \end{cases} \tag{S37}$$

The first of these two limits signifies skips so large that the position of the protein become so large that skips are uncorrelated and the scanning density is very low. In this limit, skips are indistinguishable from 3D excursions, and the search time equals the optimal search time in sliding-only regime (**Equation** S23) with $\tau_{\rm 3D}$ replaced by by $\tau_{\rm skip}$. The second of the two limits signifies when skips large but correlated, such that the effective scanning density is high even though the scanning density is low.

Having found two local optima, the more favorable search strategy is the one corresponding to the lowest search time. Hence, a combination of skipping and sliding is preferred (over just sliding) when $T^{\rm sNs}_{\rm searh} < T^{\rm sliding}_{\rm search}$. Using **Equations** S26 and S37 we in **Fig.** 3c plot the various regimes of the searcher in therms of $x_0^2$ and $y_0^2$.

#### 2.3 Skipping over the search-stability paradox

Following the reasoning of Slutsky & Mirny's original formulation of the search-stability paradox, we envision a protein to adopt two distinct conformations (**Supplementary Figure** 1d): One with strong sequence specificity and low diffusion constant $D_{\rm rec}$, and one with little specificity and high diffusion constant $D_{\rm srch}$. In the context of sNs search, the non-specific fast search mode corresponds to diffusive skip, with mean-squared skipping length is given by the diffusion constant in the fast mode and the time needed to perform a skip

$$l_{\rm skip}^2 = D_{\rm srch}\tau_{\rm skip}. \tag{S38}$$

If the protein adopts an optimal sNs search, its search time is given by (taking the lower limit of **Equation** S37)

$$T_{\rm search}^{\rm sNs} \approx 2L\sqrt{\frac{\tau_{\rm 3D}}{D_{\rm srch}}} \tag{S39}$$

Hence, the search time is dictated solely by the diffusion constant in the fast search mode, and not the diffusion constant $D_{\rm rec}$ of the recognition mode.

### References


[1] Tao Ju Cui, Misha Klein, Jorrit W Hegge, Stanley D Chandradoss, John van der Oost, Martin Depken, and Chirlmin Joo. Argonaute bypasses cellular obstacles without hindrance during target search. *Nature Communications*, 10(1):4390, 2019.

[2] Sidney Redner. *First-Passage Fundamentals.* Cambridge University Press, 2001.

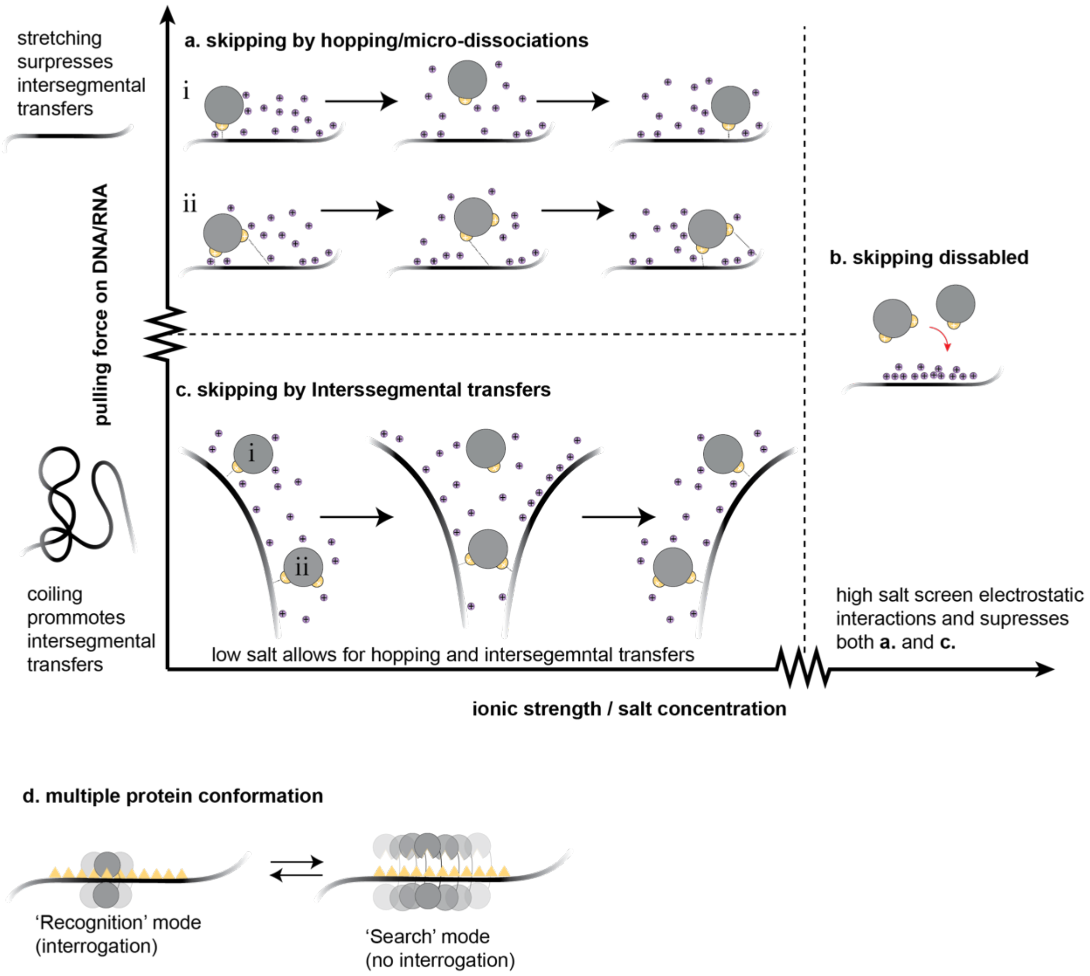


**Supplementary Figure 1| Various manifestations of skipping . a,** _Hopping /micro-dissociations:_ The protein is envisioned as transitioning between being tightly bound to (specific interactions) and loosely associated with (non-specific electrostatic interactions) the substrate. Counterions (purple) condensate upon the protein loosening it grip, the protein diffuses to a nearby (correlated) DNA site, where it attaches again after displacing the condensed counterions. Proteins with two or more binding residues (ii) can 'walk' along the substrate by alternating which residue is attached/detached. **b,** Increasing the ionic buffer strength will induce more screening, and reduce the number of hopping events that can be completed before complete dissociation. **c,** _Intersegmental transfers:_ As DNA/RNA polymers are flexible, two distant sites along the contour can come in close proximity through bending. If the protein possesses multiple binding pockets, the searcher can temporally hold on to two polymer segments before making it across. Even for proteins with a single binding pocket, intersegmental transfers can still happen when segments are brought close enough. Applying a mechanical pulling force to stretch the DNA/RNA will reduce the contribution of intersegmental transfers, and increase those of hops along segments (**a**). **d,** _Search-recognition modes:_ Particular target searchers have been shown to adopt multiple protein conformations exhibiting varying interaction strengths with the substrate. In the recognition mode, the protein interacts strongly with the DNA, thereby interrogating every sequence along its path (yellow triangles). In the search mode, specific interactions are reduced, and lateral diffusion is sped up. Although originally not envisioned as such, also this 'search mode' can be seen as a form of base skipping as during such fast lateral motion it is assumed the protein has no time to properly interrogate the underlying sequences.

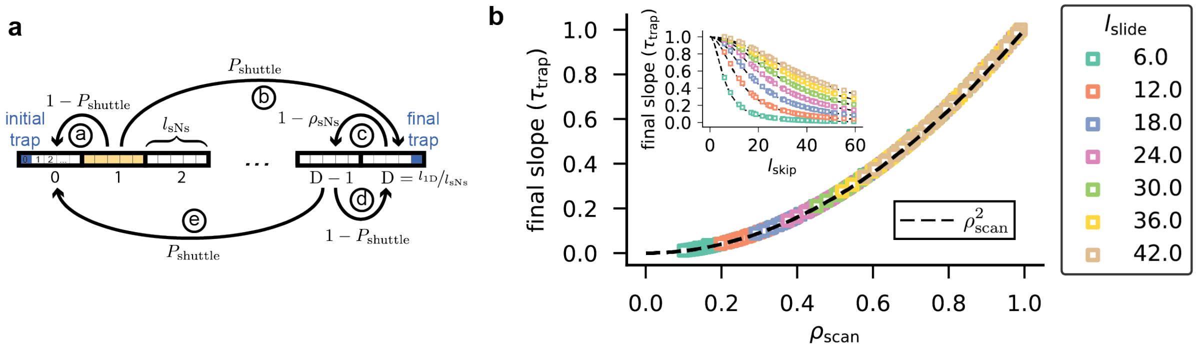


**Supplementary Figure 2| Shuttling time at large trap separations. a,** A schematic showing the possible paths to cross from one trap to the other. To calculate the probability to return to the first segment (number 0) after the first skip (segment 1 shown in yellow), without getting trapped in the second trap (rightmost blue square), we sum the probability over the possible paths through which this can be realized in the coarse-grained model. The possible paths are $a$ and $b+(c+d)^m+c+e$ for any number of cycles $m \in [0,1,2,\dots]$, and their respective probabilities are indicated next to their arrows. The original model is coarse grained to the level where each segment has the length $l_{\mathrm{sNs}}$, and thus the coarse grained index runs from $D=0$ to $D=l_{1D}/l_{\mathrm{sNs}}$. **b,** Final slope of $T_{\mathrm{shuttle}}$ (in units of the trapping time $\tau_{\mathrm{trap}}$) in our numerical calculations of the model versus scanning density. Inset shows the final slope as a function of skipping length. Different colored markers correspond to different values of the sliding length. See **Supplementary Information** for details on parameter sweep. Solid line represents our scaling arguments, and the complete data collapse visible indicates that the terminal slope of $T_{\mathrm{shuttle}}$ grows as $\rho^2_{\mathrm{sNs}}$.

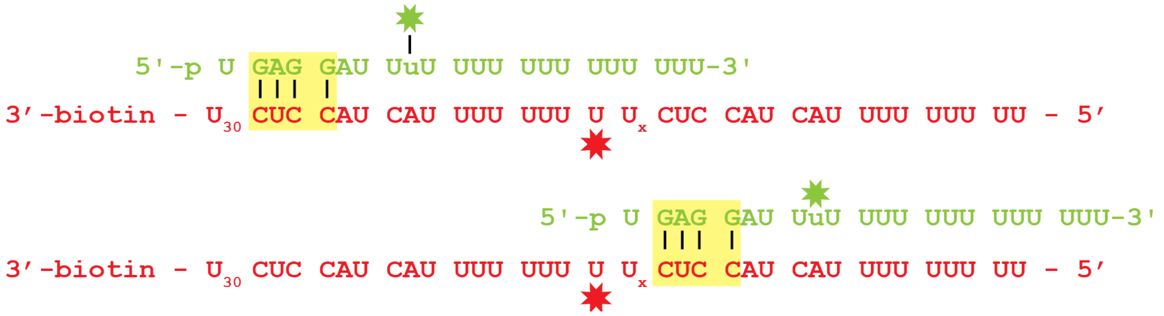


**Supplementary Figure 3| construct design hAgo2 substrate for smFRET.** ssRNA constructs (red) are passivated to the microscope slide using a 3'-biotin-streptavidin linkage. The two trapping sequences, 4 nt sequences that are complementary to the corresponding guide nucleotides (green), are highlighted in yellow. Top figure represents the ‘high FRET’ configuration, while the bottom figure displays Ago bound to the trap resulting in ‘low FRET’. The distance between traps is varied by adding uracil nucleotides (Ux reads: ‘x times a U’). To embed the traps within the sequence, as opposed to them being the outermost sites, poly-U sequences flank both traps.

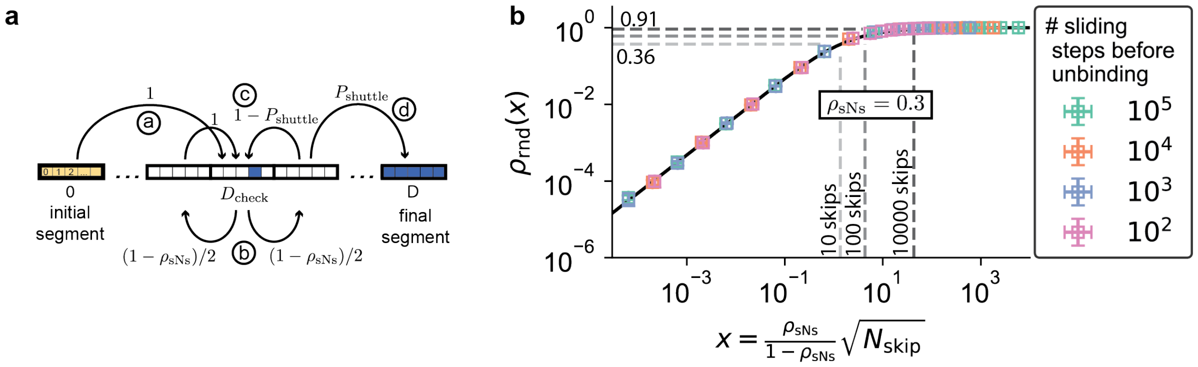


**Supplementary Figure 4| The effective scanning density of facilitated diffusion. a,** To approximate a lateral diffusion event, we assume the protein starts its search in the leftmost segment (yellow) and 'unbinds' when it reaches segment $D = l_{\text{rnd}}/l_{\text{sNs}}$ (blue rectangle). To calculate the probability of unbinding before having visited a particular site (blue square) located within segment $D_{\text{check}}$, we sum the probability of the possible paths through which this can be realized. The possible paths are $a + (b + c)^m + b + d$ for any number of cycles $m \in [0,1,2,\dots]$, and their respective probabilities are indicated next to their arrows. **b,** Comparison of $\rho_{\text{rnd}}(x)$ (solid line, **Equation S16**) to Monte Carlo simulations (symbols). Dashed lines indicate $\rho_{\text{rnd}}$ values for hAgo2's ($\rho_{\text{sNs}} \approx 0.3$) that typically skip 10 (lightest grey), 100 (middle grey), and 1000 times (dark grey) before unbinding.

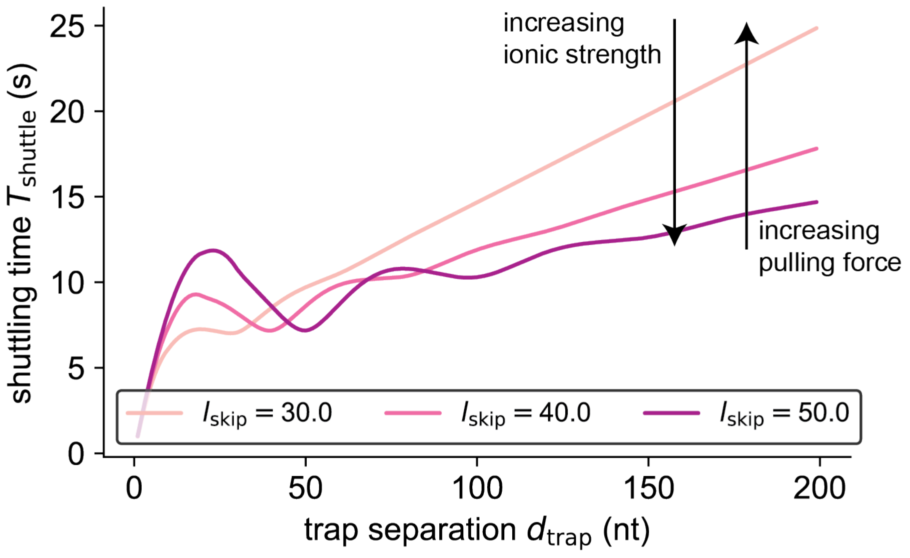


**Supplementary Figure 5| The effect of varying skipping length.** Shuttling time vs. trap separation for $l_{\text{slide}} = 10$ and increasing $l_{\text{skip}}$, mimicking the effect of either increasing salt concentration or decreasing the pulling forces on the substrate.
